# Multimodal immersive trail making – virtual reality paradigm to study cognitive-motor interactions

**DOI:** 10.1101/2020.05.27.118760

**Authors:** Meir Plotnik, Oran Ben-Gal, Glen M. Doniger, Amihai Gottlieb, Yotam Bahat, Maya Cohen, Shani Kimel-Naor, Gabi Zeilig, Michal Schnaider Beeri

**Affiliations:** Center of Advanced Technologies in Rehabilitation, Sheba Medical Center, Ramat Gan, Israel; Department of Physiology and Pharmacology, Sackler Faculty of Medicine, Tel Aviv University, Tel Aviv, Israel; Sagol School of Neuroscience, Tel Aviv University, Tel Aviv Israel; Joseph Sagol Neuroscience Center, Sheba Medical Center, Ramat Gan, Israel; Department of Neurological Rehabilitation, Sheba Medical Center, Ramat Gan, Israel; Deartment of Rehabilitation, Sackler Faculty of Medicine, Tel Aviv University, Israel; Department of Psychiatry, The Icahn School of Medicine at Mount Sinai, New York, NY USA

**Keywords:** Executive functions, Cognitive-motor interactions, Construct validity, Divided attention, Neuropsychological testing, Virtual reality

## Abstract

**Background:** Neuropsychological tests of executive function have limited real-world predictive and functional relevance. An emerging solution for this limitation is to adapt the tests for implementation in virtual reality (VR). We thus developed two VR-based versions of the classic Color-Trails Test (CTT), a well-validated pencil-and-paper executive function test assessing sustained (Trails A) and divided (Trails B) attention - one for a large-scale VR system (DOME-CTT) and the other for a portable head-mount display VR system (HMD-CTT). We then evaluated construct validity, test-retest reliability, and age-related discriminant validity of the VR-based versions and explored effects on motor function.

**Methods:** Healthy adults (*n*=147) in three age groups (young: *n*=50; middle-aged: *n*=80; older: *n*=17) participated. All participants were administered the original CTT, some completing the DOME-CTT (14 young, 29 middle-aged) and the rest completing the HMD-CTT. Primary outcomes were Trails A and B completion times (t_A_, t_B_). Spatiotemporal characteristics of upper-limb reaching movements during VR test performance were reconstructed from motion capture data. Statistics included correlations and repeated measures analysis of variance.

**Results:** Construct validity was substantiated by moderate correlations between the ‘gold standard’ pencil-and-paper CTT and the VR adaptations (DOME-CTT: t_A_ 0.58, t_B_ 0.71; HMD-CTT: t_A_ 0.62, t_B_ 0.69). VR versions showed relatively high test-retest reliability (intraclass correlation; VR: t_A_ 0.60-0.75, t_B_ 0.59-0.89; original: t_A_ 0.75-0.85, t_B_ 0.77-0.80) and discriminant validity (area under the curve; VR: t_A_ 0.70-0.92, t_B_ 0.71-0.92; original: t_A_ 0.73-0.95, t_B_ 0.77-0.95). VR completion times were longer than for the original pencil-and-paper test; completion times were longer with advanced age. Compared with Trails A, Trails B target-to-target VR hand trajectories were characterized by delayed, more erratic acceleration and deceleration, consistent with the greater executive function demands of divided vs. sustained attention; acceleration onset later for older participants.

**Conclusions:** The present study demonstrates the feasibility and validity of converting a neuropsychological test from two-dimensional pencil-and-paper to three-dimensional VR-based format while preserving core neuropsychological task features. Findings on the spatiotemporal morphology of motor planning/execution during the cognitive tasks may lead to multimodal analysis methods that enrich the ecological validity of VR-based neuropsychological testing, representing a novel paradigm for studying cognitive-motor interactions.

## Background

The term “executive functions” is an umbrella term for a wide range of cognitive processes and behavioral competencies necessary for the cognitive control of behavior, for example, problem solving, planning, sequencing, the ability to sustain attention, utilization of feedback, multitasking etc., (1). Neuropsychological tests of executive functions aim to assess these processes (2). Accordingly, performance on these tests is assumed indicative of executive functioning in everyday living (3). One of the limitations of these tests relates to their low ‘ecological validity’, i.e. the uncertainty about how closely they reflect an individuals’ executive functions capacity in real life (4-6). In this regard, Burgess et al. (7) claimed that “the majority of neuropsychological assessments currently in use were developed to assess ‘cognitive constructs’ without regard for their ability to predict ‘functional behavior’”.

### Neuropsychological assessment in Virtual Realtiy (VR)

Early discussions of ecological validity in neuropsychology emphasized that the technologies available at that time could not replicate the environment in which the behavior of interest would ultimately take place (8). Furthermore, currently, most neuropsychological assessments still use outdated methods (e.g., pencil-and-paper assessments; static stimuli) that have yet to be validated with respect to real-world functioning (9).

To overcome this limitation, testing participants in real word situations (e.g., the Multiple Errands Test-MET (10)) was considered as ecologically valid and advantageous alternative to traditional tests (11). However, this approach have some logistically related drawbacks (e.g., travel to a testing site) (12).

In an attempt to address these drawbacks, the Virtual Errands Test (VET) was devised by McGeorge et al. (13) to adapt the MET for VR - based administration. Yet, this test, and similar VR variants, are limited in their ability to distinguish between healthy and clinical cohorts (see (11) for a review) and to invoke performance in the virtual tasks similar as performance in the real world (e.g., (14, 15)). Further, most of these VET like and other VR based – tests use a simulated VR environment presented on a standard computer screen (e.g., Elkind et al., (16)), which may lead to lack of immersion and thus paradoxically exacerbate rather than ameliorate ecological validity.

On the other hand, some VR environments simulating shopping tasks for the assessment of executive function demonstrated good ecological validity (17, 18). Notwithstanding these encouraging findings, the approach of adapting executive function testing for the VR environment has not yet been widely accepted in both research and clinical contexts.

### Research rational

Critically, we posit that the concept of ‘ecological validity’ is not merely related to the type of task performed and its relevance to daily living. In general, each response on a cognitive task involves interactions with sensory and motor functions, first in order to determine the type of response needed and then to plan and execute the appropriate behavioral response. These processes cannot be separated and examined with traditional pencil-and-paper testing or even with computerized testing platforms.

Thus, as a first stage, we aim to develop VR neuropsychological tests by adapting an existing well-validated traditional neuropsychological test that measure specific cognitive constructs, i.e., keeping the focus on the core cognitive domain. These adaptations will enhance the ecological validity by also including multi-multimodal (e.g., cognitive-sensory-motor) interactions. These VR-based adaptations may allow measurement of cognitive function in a manner more relevant to performance of daily life activities, including the interaction among multiple functions (19-24). In particular, VR technology facilitates collection of quantitative three-dimensional kinematic data (unavailable for traditional neuropsychological tests) that tracks motion in space and may provide greater sensitivity (i.e., ability to discriminate and define levels of performance).

### The Color Trails Test (CTT)

The Trail Making Test (TMT) (25, 26) is among the most popular pencil-and-paper tests of executive function, attention and processing speed in research and clinical neuropsychological assessment. The Color Trails Test (CTT) is a culture-fair variant of the TMT. In *Trails A* the participant draws lines to sequentially connect circles numbered 1 to 25 (odd numbered circles are pink, even numbered circles are yellow). In *Trials B* participant alternates between circles of two different colors (i.e. 1-pink, 2-yellow, 3-pink, 4-yellow, etc.) (27). Scoring is based on the time needed to complete the tasks, with shorter time indicating better performance. It has been proposed that *Trails A* assesses sustained visual attention involving perceptual tracking and simple sequencing, and Trails B more directly assesses executive function, including such sub-processes as divided attention, simultaneous alternating and sequencing (27, 28).

### The present study

The overall goal of this project was to examine the value of translating of a well-validated executive functions task currently administered using pencil and paper to a VR testing paradigm. We developed two VR adaptations of the CTT test – the DOME-CTT, designed for a large-scale VR system in which the participant stands in the middle of a visual scene projected on a 360° dome-shaped screen, and the HMD-CTT, designed for a low-cost head-mount device (HMD), in which the tasks are presented to the participant via VR goggles. In addition to developing the VR-based tests, we evaluated their ability to measure the same cognitive constructs (construct validity) as the (gold standard) pencil-and-paper CTT, as well as their ability to differentiate among healthy young, middle-aged and older age groups (discriminant validity) compared with the original CTT. Finally, we explored cognitive-motor interactions during performance of the VR-CTT tasks.

## Methods

### General

Two platforms of VR-CTT were developed: DOME-CTT and HMD-CTT. Findings from experiments performed with these platforms are described in *Study 1* and *Study 2*, respectively. There were a total of 147 healthy participants in Study 1 and Study 2 (within the framework of larger, different experimental protocols - see *table S1*, in *supplementary file #1*) subdivided into the following age groups: (1) young adults (YA), ages 18–40 years (*n*=50); (2) middle-aged adults (MA) ages 40-65 years (*n*=70); (3) older adults (OLD), ages 65-90 years (*n*=17). For all groups, exclusion criteria were motor, balance, psychiatric or cognitive conditions that may interfere with understanding the instructions or completing the required tasks (determined based on screening interviews). The protocols were approved by the Sheba Medical Center institutional review board (IRB), and all participants signed informed consent prior to entering the study.

### Methods for Study 1 (DOME-CTT)

#### Participants

Data from 14 YA (age: 27.9±5.0 [mean±SD] years, education: 16.4±2.9 [mean±SD] years; 9 females) and 29 MA (age: 55.8±6.2 years, education: 16.3±3.0 years; 16 females) were included in Study 1.

#### Apparatus

A fully immersive virtual reality system (CAREN High End, Motek Medical, The Netherlands) projected a virtual environment consisting of the task stimuli on a full-room dome-shaped screen (Fig. 1). The system comprises a platform with an embedded treadmill, and is synchronized to a motion capture system (Vicon, Oxford, UK). Auditory stimuli and feedback are delivered via a surround sound system.

**Figure 1:**
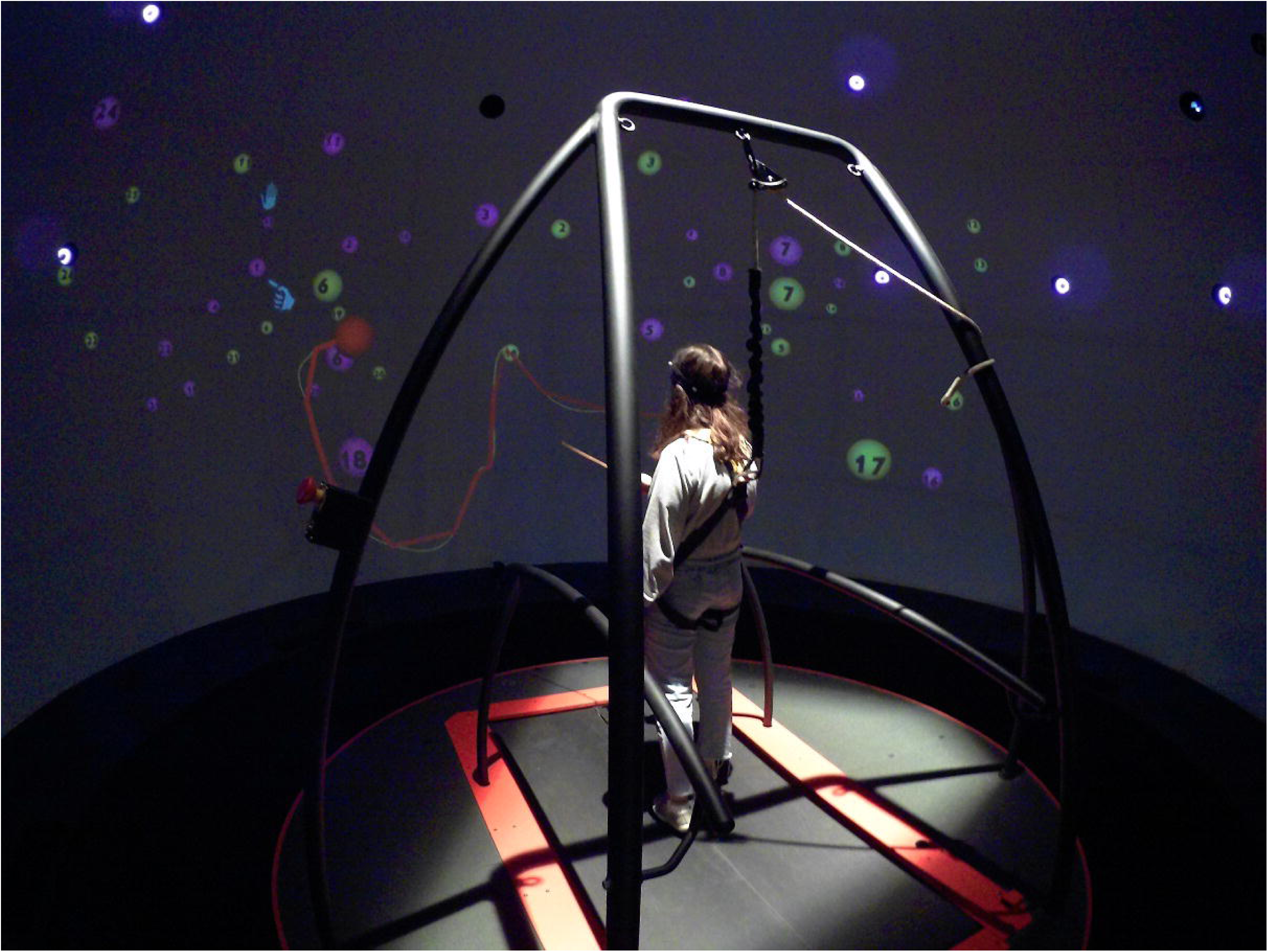
The CAREN High End (Motek Medical, The Netherlands) has a 6-degree of freedom (three translation axes, three rotation axes) moveable platform that is synchronized with a virtual visual scene projected on a 360° dome-shaped screen. A split-belt treadmill is embedded in the platform (not operated in this study). A motion capture system (Vicon) and a pair of force plates embedded in the treadmill provide data on the participant’s spatiotemporal position during the task (sampling rate 120Hz). The participant who performs the DOME-CTT VR adaptation (see text), is secured with safety harness, holding a wand-like pointing stick with a marker on its edge in order to control the avataric red ball and to move it towards the target-balls.

#### Adapting the pencil-and-paper Color Trails Test to a large-scale VR system (DOME)-based test – The DOME-CTT (Figure 2)

A virtual version of the CTT was developed to demonstrate the feasibility of performing neuropsychological testing in a virtual environment. The original pencil-and-paper CTT consists of four parts: practice (Trails) A, test (Trails) A, practice (Trails) B and test (Trails) B (27). As below, all were adapted to the VR environment. In the VR version of the CTT, the two-dimensional (2D) page (Fig. 2A) is replaced with a three-dimensional (3D) VR space (Fig. 2B, C) that introduces a new depth dimension to the target balls (i.e., replacing the 2D circles) and to the generated trajectory. The translation to 3D geometrics followed principals governing the 2D design (compare Fig.2A and Figs. 2B, C). For example: (1) balls were placed so that virtual trajectories between two balls would not cross previously generated trajectories (i.e., between pairs of balls to be touched by the participant earlier in the task sequence); (2) proximity of balls in any given portion of the 3D space was similar to that of the 2D space in the original CTT; (3) for Trails B, we position the corresponding ball of the incorrect color at a similar relative distance to the correct ball to the original 2D CTT.

**Figure 2:**
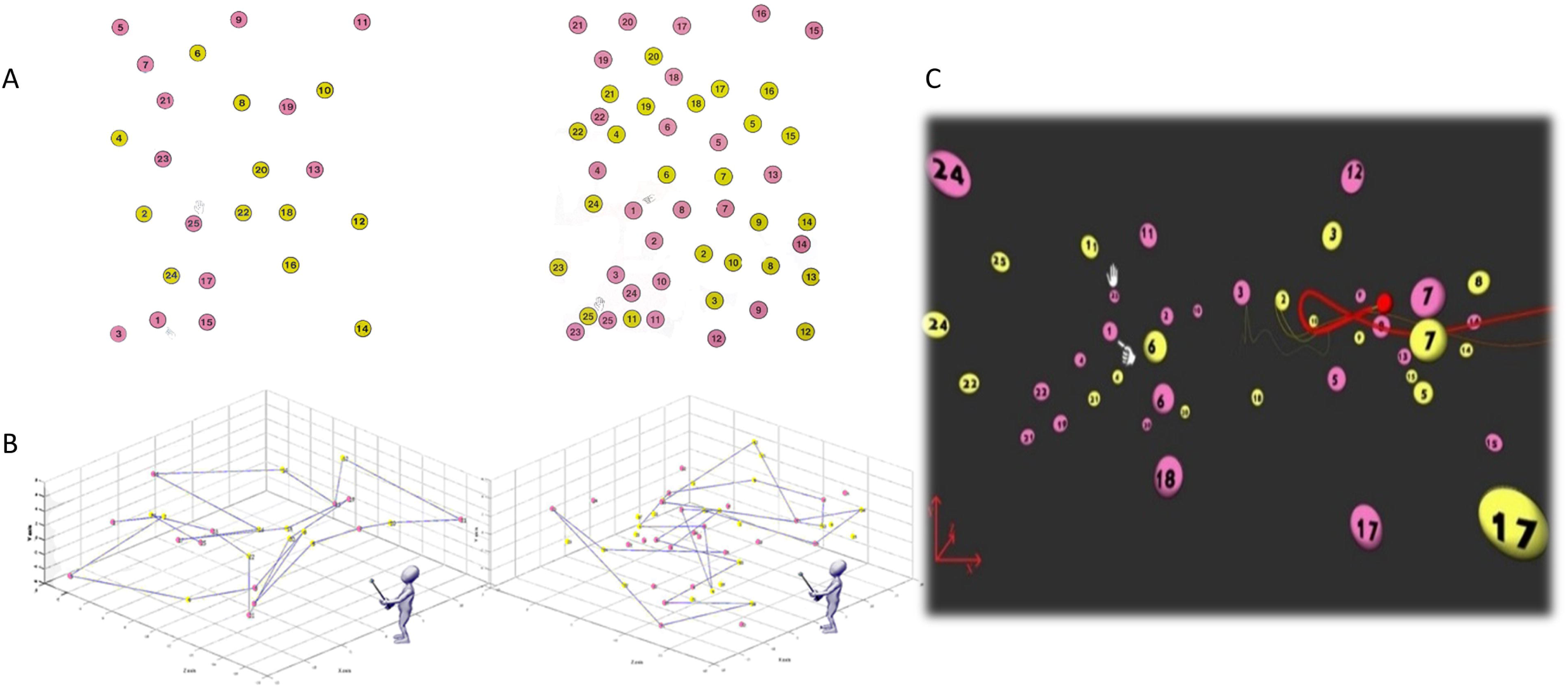
Adaptation of the pencil-and-paper CTT to yield the DOME-CTT. **A**. Trails A (left) and B (right) tasks from the pencil-and-paper CTT. The first and last balls are each indicated by a picture of a hand (pointing and stop signing, respectively, seen here in Trails B only). **B**. Schematic depiction of the spatial geometric orgnizaiton of the target balls in the three dimensional space. **C**. Trails B task from the DOME-CTT, as viewed by the participant. Ball size varies as a function of its proximity to the participant in the VR space. A fixed pointing stick marker is represented by the red ball. Its trace is depicted in yellow, while the red, more pronounced trace is used for visual feedback in the VR space. The first and last balls are indicated using the hand icon.

The participant performed the DOME-CTT with a marker affixed to the tip of a short wand-like pointing stick held in the dominant hand (corresponding to the pen or pencil in the original CTT). The three-dimensional coordinates of the marker are tracked in real time by the motion capture system at a sampling rate of 120 Hz. A virtual representation of this marker appears within the visual scene (i.e., ‘avatar’, represented by a small red ball – Fig. 2C). To mimic drawing lines in the 2D pencil-and-paper CTT, as the participant moved his/her hand within the VR space, a thick red ‘tail’ trailed directly behind the position of the (red ball) avatar, gradually becoming a faint yellow tail as the avatar is moved further away from the initial position (Fig. 2C).

Movement of the marker was recorded in real time by a motion capture system that allows the reconstruction of kinematic data over the entire test duration.

In addition to the apparatus, the testing procedure was also adapted for the new format. As above, the original pencil-and-paper CTT comprises four consecutively administered test levels: (1) Trails A practice; (2) Trails A; (3) Trails B practice; and (4) Trails B [16]. Though drawing lines with a pen/pencil on a piece of paper is highly familiar, manipulation of the VR ‘controller’ (i.e., the marker affixed to the pointing stick) to move an avatar (i.e., the red ball) within the virtual environment is a relatively unfamiliar skill. Thus, the DOME-CTT began with an additional practice level in which participants practiced guided movement of the avatar within the virtual space to touch numbered ball targets. During this level, participants were introduced to the positive feedback received when touching the correct ball (i.e. momentary enlargement of the ball) and the negative feedback upon touching an incorrect ball (i.e., brief buzzing sound). After this initial practice level, test levels corresponding to those in the original CTT were administered. However, unlike the pencil-and-paper CTT, Trails A and Trails B were each preceded by two different practice levels. In the first practice level, all virtual balls were clustered near the center of the visual field, and in the second practice level, the balls were spread throughout the visual field, approximating the spatial distribution of the balls in the actual testing levels. A video demonstration of the DOME-CTT is provided in *supplementary file* #2.

#### Procedure

Data on pencil-and-paper CTT and DOME-CTT were collected as part of three different experimental protocols (see *table S1* in *supplementary material file #1*). All data (with the exception of test retest data) described in this study were collected on the first visit. The participants completed the pencil-and-paper CTT and DOME-CTT on the same day in counterbalanced order across participants.

#### Outcome measures and statistical analysis

For the pencil-and-paper CTT and the DOME-CTT, completion times for Trails A and B were recorded (t_A_, t_B,_ respectively). Construct validity was assessed by correlating t_A_ and t_B_ from the DOME-CTT with the corresponding scores from the gold standard CTT (Pearson coefficient). Analysis of variance (ANOVA) was used to assess effects of Group (young, middle aged; between-subjects factor), Trails (A and B; within -subjects factor) and Format (pencil-and-paper CTT, DOME-CTT; within-subjects factor). Partial Eta Squared was computed as a measure of effect size. To verify suitability of parametric statistics, Shapiro-Wilk normality tests were run for each outcome variable per group. Of the eight normality tests, none indicated non-normal distributions (Shapiro-Wilk statistic ≤ .93; p ≥ .16). Summary statistics (mean ± SD) were computed for t_A_ and t_B_, recorded in the paper and pencil CTT and DOME-CTT.

Errors were manually recorded by the experimenter for the pencil-and-paper CTT (27) and both manually and automatically by the DOME-CTT software. Related samples Wilcoxon Sign Test (non-parametric) test was used to evaluate Format effect separately for Trails A and B. Mann-Whitney U tests were used to evaluate group effect.

To evaluate discriminant validity (i.e., ability to separate between YA and MA) of the DOME-CTT, as compared with the pencil-and-paper CTT, we plotted receiver operating characteristic curves (ROC) for Trails A and Trails B (i.e., t_A_ and t_B,_ respectively) for each test version and calculated the area under the curve (AUC; range: 0-1, higher values reflect better discriminability).

Level of statistical significance was set at 0.05. Statistical analyses were run using SPSS software (SPSS Ver. 24, IBM).

### Methods for Study 2

#### Participants

Data from 36 YA (age: 26.7±4.1 [mean±SD] years, education: 15.9±2.3 [mean±SD]; 21 females), 51 MA (age: 56.2±6.2 years, education: 16.8±3.0 years; 39 females) and 17 OLD (age: 73.7±6.5 years, education: 13.1±2.7 years; 11 females) were included in Study 2.

#### Apparatus

VR technologies have advanced extremely rapidly in recent years. In addition to new technical features for precise stimulus delivery and response measurement, as well as enhanced usability, low-cost VR is now widely accessible. The most accessible type of VR system is the head-mount device (e.g., HTC Vive, Oculus Rift), which is designed for home-based operation. In addition to its continued popularity for entertainment, VR is now being applied in a variety of ‘serious’ contexts, ranging from surgical simulation to the study of human performance and psychological function (29, 30). For this study, we used a fully immersive VR system (HTC; New Taipei City, Taiwan) including a headset with ∼100° field of view (FOV) angle in the horizontal plan and ∼110° FOV angle in the vertical plan. Also included were a controller for user interaction with the virtual environment and two *‘*lighthouse*’* motion trackers for synchronizing between actual controller position and corresponding position in the virtual environment.

#### Adapting the pencil-and-paper Color Trails Test to a headset-based VR system – The HMD-CTT

In developing the HMD-CTT version, we adopted a similar approach to the development of the DOME-CTT but for a VR headset system. Briefly, we used the popular Unity3D VR game engine (31) to design a virtual environment for the CTT. With the exception that the participant held the HTC controller rather than a wand-like pointing stick, task design paralleled the DOME-CTT, including positive and negative feedback cues, practice and test procedures. Figure 3 illustrates how the 2D format of the original CTT was translated to the 3D HMD-CTT format (Trails A; see also video demonstration in *supplementary file* #3). Similarly to training phases for the DOME-CTT described above, pre-test training and acclimatization trials were prepared and performed as part of the HMD-CTT procedure.

**Figure 3:**
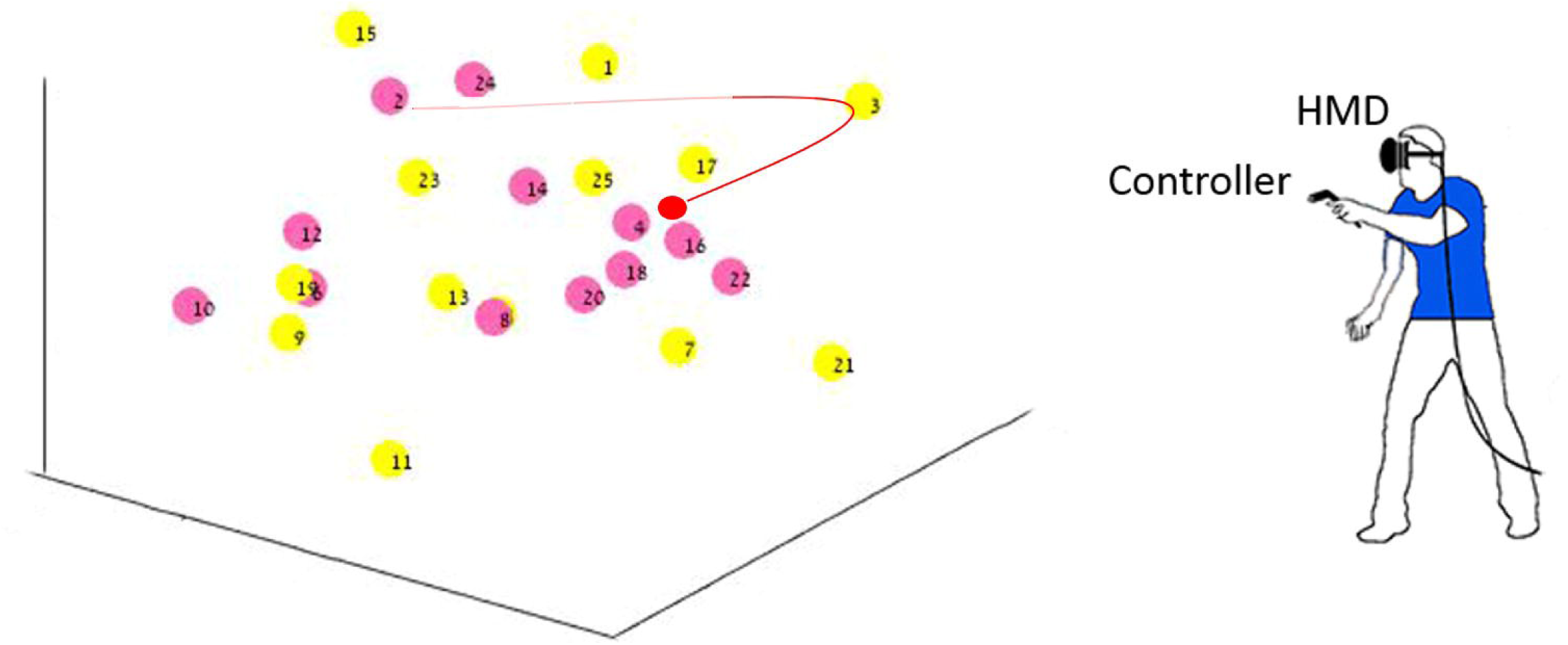
Translation of pencil-and-paper CTT to HMD-based VR format. In the three-dimensional VR-based HMD-CTT, the participant completes the task while standing and wearing a VR headset. S/he holds a VR controller in his/her dominant hand to move the (red ball) avatar to sequentially connect numbered balls distributed throughout the virtual space visible in the headset. (Trails A shown.)

#### Procedure

The procedure was identical to that of *Study 1* (see above).

#### Outcome measures and statistical analyses

For the pencil-and-paper CTT and the HMD-CTT, completion times for Trails A and B were recorded (t_A_, t_B,_ respectively). As for *Study 1*, construct validity of t_A_ and t_B_ was assessed by correlating t_A_ and t_B_ from the HMD-CTT with the corresponding scores from the gold standard pencil-and-paper CTT. Similar to the DOME-CTT study, repeated-measures ANOVA was used to assess the effects of Group (YA, MA, OLD; between-subjects factor), Trails (A and B; within -subjects factor) and Format (pencil-and-paper CTT, HMD-CTT; within-subjects factor). Partial Eta Squared was computed as a measure of effect size. We used the Bonferroni correction for adjustment for multiple comparisons in the post-hoc pairwise comparisons.

As for *Study 1*, Shapiro-Wilk normality tests were run for each outcome variable per group to verify the suitability of parametric statistics. Of the twelve normality tests, four indicated non-normal distributions: t_A_ pencil-and-paper CTT, MA group, Shapiro-Wilk statistic=.786, p=.002; t_B_ pencil-and-paper CTT, MA group, Shapiro-Wilk statistic=.770, p=.002; t_B_ pencil-and-paper CTT, YA group, Shapiro-Wilk statistic=.744, p=.001, t_A_ pencil-and-paper CTT, OLD group, Shapiro-Wilk statistic=.844, p=.015. Additional analyses used are mentioned in the *Results* section.

Regarding prevalence of errors, related samples Wilcoxon Sign Test (non-parametric) test was used to evaluate Format effect separately for Trails A and B. Kruskal-Wallis tests was used to evaluate group effect (three levels) separately for Trails and Format.

As for *Study 1*, level of statistical significance was 0.05, and statistical analyses were run with SPSS.

#### Qualitative analysis of manual performance

Spatial coordinates of the controller position (corresponding to ‘red ball’ avatar) were recorded throughout the HMD-CTT sessions. Custom software written in MATLAB (Mathworks, Inc.) used this data to extract and analyze the 24 target-to-target reaching movements during Trails A and Trails B, respectively (errors were excluded). We performed qualitative assessment of the trajectories generated by participants in each of the three groups by examining the grand averages of their velocity profiles in an attempt to characterize upper-limb motor behavior associated with performance of the HMD-CTT tasks. For a full description of the methodology used to generate these grand averages see *supplementary file #1*.

### Evaluation of Test-retest Reliability (Study 1 and Study 2)

To evaluate test-retest reliability, some participants completed a second assessment.

Fifteen MA participants from *Study 1* completed an additional evaluation about 12 weeks after the initial evaluation (protocol 1 described in *table S1* in *supplementary material* file #1), in which they were administered the original pencil-and-paper CTT and DOME-CTT in the same order as in the initial evaluation.

Thirty-two MA participants from *Study 2* completed an additional evaluation about 12 weeks after the initial evaluation.

Also from *Study 2*, and twenty participants (*n*=10 YA, *n*=1 MA, *n*=9 OLD) completed an additional evaluation 2 weeks after the initial one. All participants were administered the original pencil-and-paper CTT and HMD-CTT in the same order as in the initial evaluation.

To assess test-retest reliability, we computed intraclass correlation coefficients (ICC; two-way mixed, effects, absolute agreement, (32)) for t_A_ and t_B_ scores from the traditional pencil-and-paper CTT and the DOME-CTT (*Study 1*) or HMD-CTT (*Study 2*) collected at two visits. ICC reflects similarity of the obtained scores irrespective of the level of performance, reflecting not only correlation, but also agreement between measurements (33, 34). By convention ICC >.75 is considered good reliability.

## RESULTS

### Study 1

#### Performance on the DOME-CTT: Group, Trails and Format effects

All the participants completed the tests in both formats. Acclimatization periods to the DOME-CTT environment varied between participants, but usually did not exceeded 10-15 min. Due to technical malfunction data on DOME-CTT Trails A from one YA participant was not recorded.

Statistical analysis revealed effects of Group (F_1,40_ = 19.2, p<.001, η^2^=.32; longer completion time for middle-aged), large effects of Trails (F_1,40_ = 108.6, p<.001, η^2^=.73; longer completion time for Trail B) and large effects of Format (t_A_: F_1,41_ = 266.4, p<.001, η^2^=.86; longer completion time for DOME-CTT). The Group X Format and the Format X Trails interactions were also found significant (F_1,40_ = 11.8, p=.001, η^2^=.22, F_1,40_ = 29.4, p<.001; η^2^=.01, respectively). These interactions reflect a larger group effect for the DOME-CTT (i.e., as compared to the group effect for the pencil-and-paper CTT), and larger difference between trail B and A for the DOME-CTT (i.e., as compared to the pencil-and-paper CTT. None of the other interactions (i.e., Group X Trails X Format and Group X Trails) were not found statistically significant (p≥.319).

Regarding prevalence of errors, more errors occurred during performance of the DOME-CTT (Figure 4) for both Trails A and B (p<.001). Group effect was not found statistically significant (p≥.056). More errors were performed by the MA group as compared the YA group, across all Trails and Formats (p≤.002)

**Figure 4:**
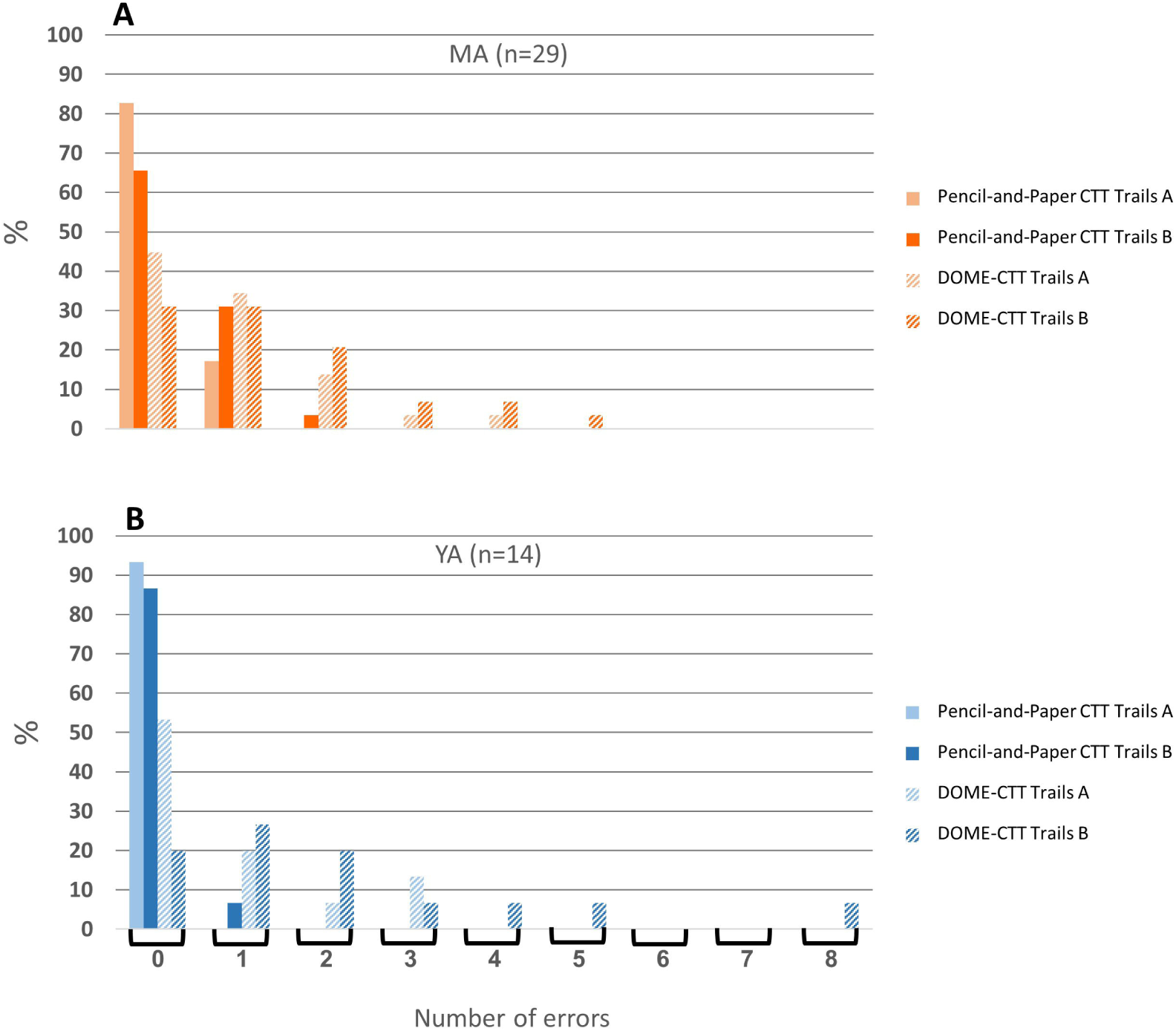
Comparison of error rates between pencil-and-paper CTT and DOME-CTT VR-based adaptation. Percent of participants making 1, 2, 3, 4, 5, 6, 7, or 8 errors for paper-and-pencil and DOME-CTT versions of Trails A and Trails B tasks (see key), separately for the MA group (panel A) and the YA group (panel B). For the pencil-and-paper CTT, most participants in both groups made no errors, and none made more than 2 errors. Conversely, for the DOME-CTT, less than half of the participants (YA and MA combined) made no errors and some made a substantial number of errors.

#### Correlations between pencil-and-paper and DOME-CTT completion times

Figure 5 shows the relationship between performance on the pencil-and-paper and DOME-CTT for the YA (blue) and MA (orange) groups. The Spearman correlation (rho; *r*_*s*_) between Part A completion time (t_A_) on the gold-standard pencil-and-paper CTT and the corresponding Part A completion time on the DOME-CTT was 0.58 (*p*<.001; Fig. 5A). For Part B completion time (t_B_), the Spearman correlation was 0.70 (*p*<.001; Fig. 5B).

**Figure 5:**
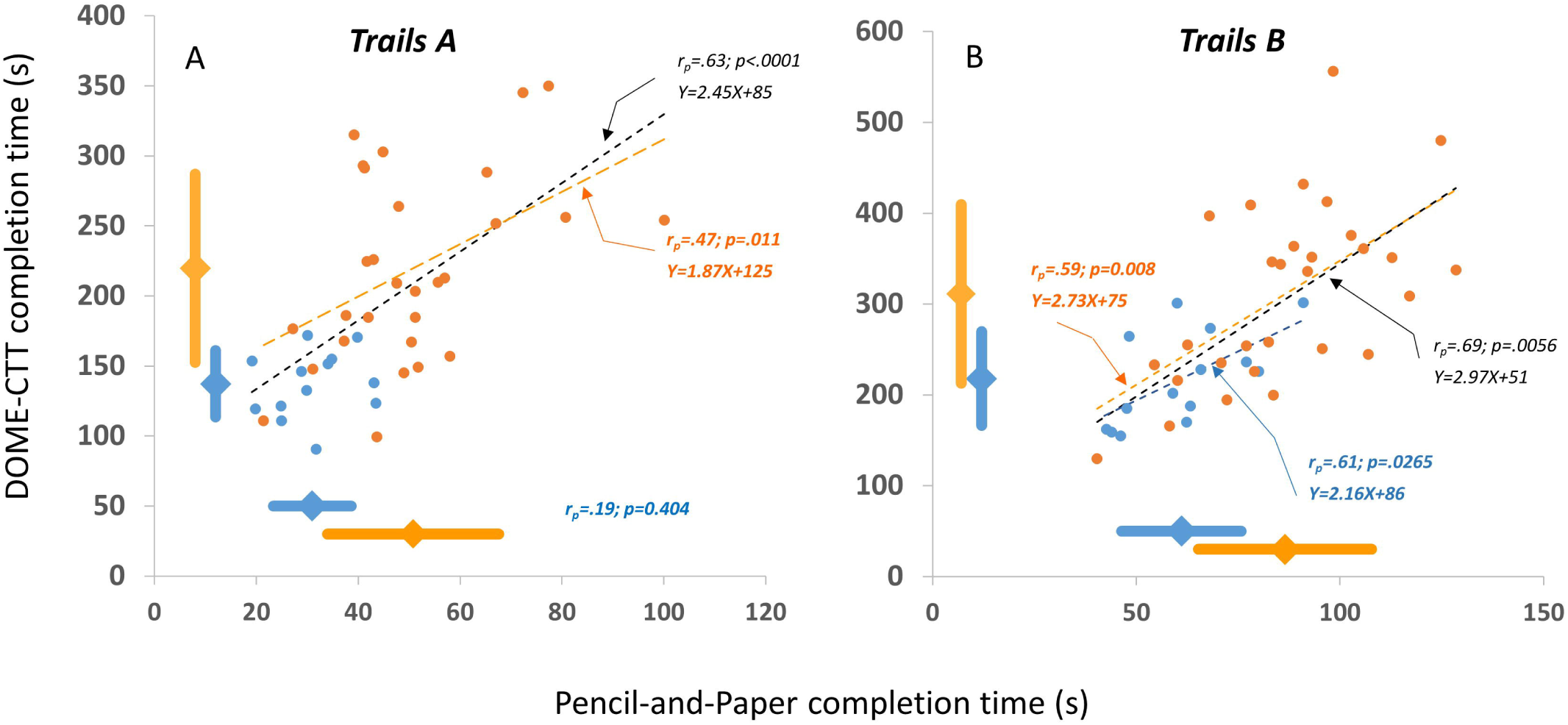
Convergent construct validity of the VR-based DOME-CTT. Trails A (*t*_*A*_, left panel) and Trails B (*t*_*B*_, right panel) completion time recorded during the gold-standard pencil-and-paper CTT plotted against the corresponding completion times recorded during the DOME-CTT for YA (blue) and MA (orange) participants. One YA datapoint is missing for Trails A, as the participant did not complete the DOME-CTT (i.e, due to technical malfunciton). Pearson correlations are shown for YA, MA, as well as combined (black) groups, and regression lines are plotted for significant correlations. Diamond markers and thick lines adjacent to the axes indicate mean ± SD for YA and MA groups.

### Study 2

#### Performance on the HMD-CTT: Group, Trails and Format effects

One OLD participant (age: 70 years, education: 12 years, female) could not perform the HMD-CTT practice and actual test levels, showing general disorientation. Another participant from this group (age: 89 years, education: 12 years, male) asked to stop the HMD-CTT during the Trails B part, expressing frustration regarding his perception of poor performance (see also Fig. 7 legend). Data from these participants were removed from related analyses.

**Figure 6:**
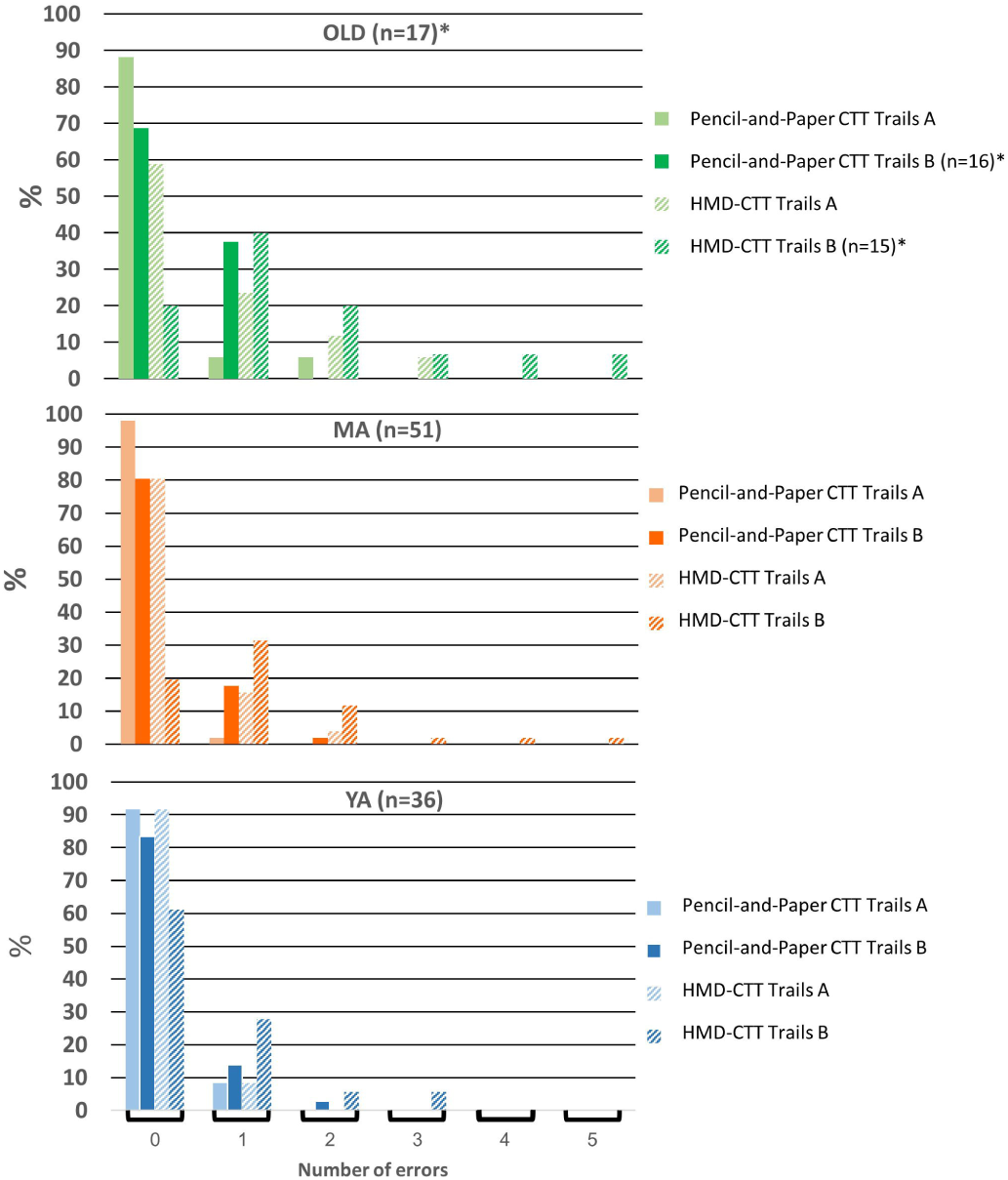
Comparison of error rates between pencil-and-paper CTT and HMD-CTT VR-based adaptation. Percent of participnats making 1, 2, 3, 4, or 5 errors for paper-and-pencil and HMD-CTT versions of Trails A and Trails B tasks (see key), separately for OLD (panel A), MA (panel B), and YA (panel C) groups. Similar to the DOME-CTT (Fig. 5), substantially more errors were made for the VR-based HMD-CTT than for the paper-and-pencil CTT; this tendency was particularly evident for OLD participants.

**Figure 7:**
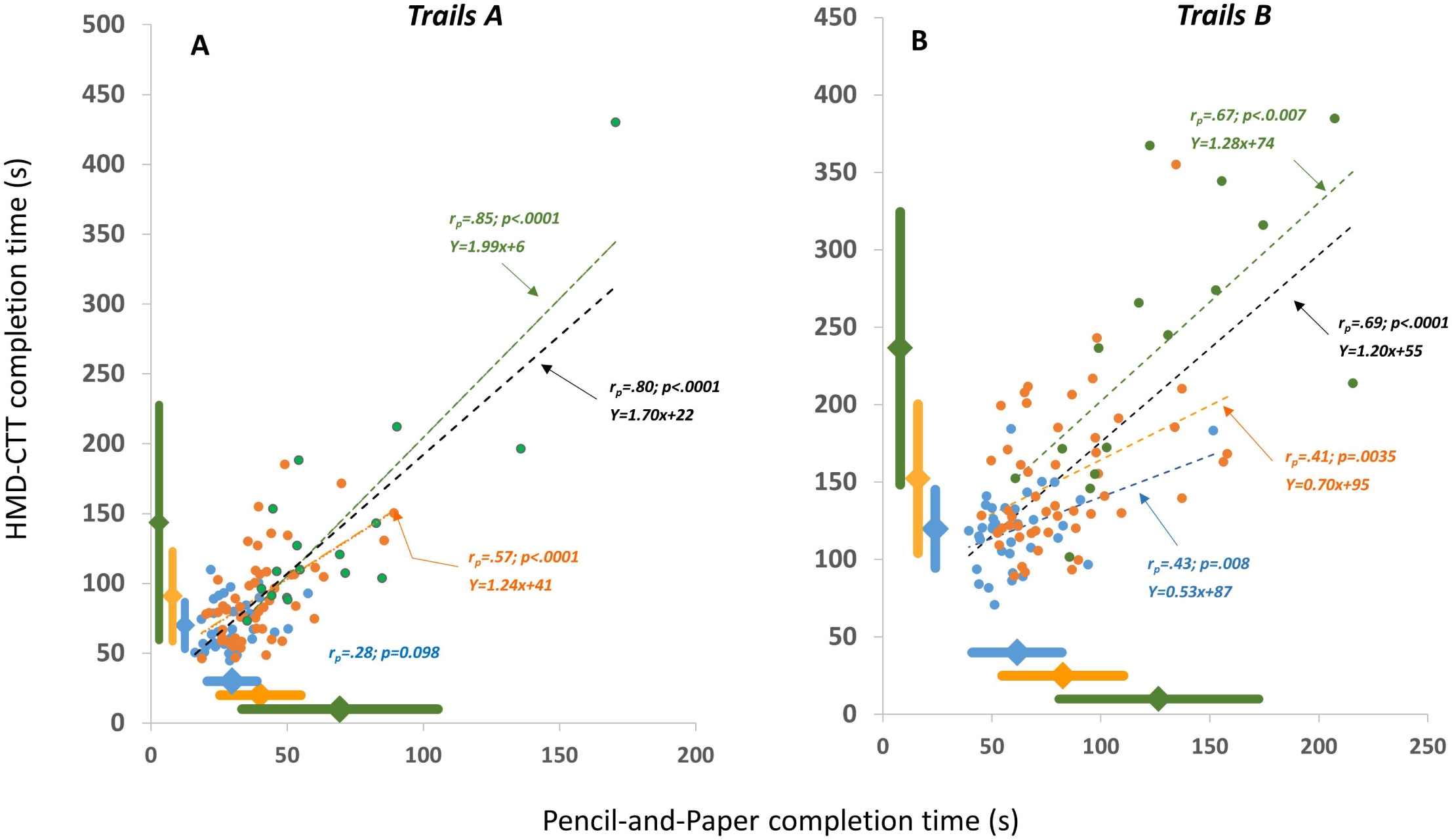
Convergent construct validity of the VR-based HMD-CTT. Trails A (*t*_*A*_, left panel) and Trails B (*t*_*B*_, right panel) completion time recorded during the gold-standard pencil-and-paper CTT plotted against the corresponding completion times recorded during HMD-CTT, for YA (blue), MA (orange), and OLD (green) participants. Two OLD datapoints are missing as the participants did not complete the HMD-CTT.* Pearson correlations are shown for YA, MA, OLD and combined (black) groups, and regression lines are plotted for significant correlations. Dimond markers and thick lines adjacent to the axes indicate mean ± SD for YA, MA, and OLD groups. *One OLD participant who could not finish the HMD-CTT Trails B had the slowest Trails A completion time, for both pencil-and-paper (i.e., 170.5 s) and HMD versions (450.2 s). His completion time for the pencil-and-paper Trails B was 327.3 s (this data point was not plotted).

Statistical analysis revealed significant effects of Group (F_2,98_ = 40.1, p<.001, η^2^=.45; progressively longer completion time with more advanced age, all post-hoc pairwise comparisons p<.001), Trails (F_1,98_ = 357.0, p<.001, η^2^=.78; longer completion time for Trail B) and Format (F_1,98_= 455.2, p<.001, η^2^=.82; longer completion time for HMD-CTT). The Group x Format, Group X Trails and Format X Trails interactions were also statistically significant (F_2,98_ = 9.9, p<.001, η^2^=.16, F_2,98_ = 14.1, p<.001, η^2^=.22, F_2,98_ = 30.3, p<.001, η^2^=.23, respectively). These interactions seem to be the results of a larger group effect for the HMD-CTT vs. the pencil-and-paper CTT, a larger group effect for Trails B vs. Trails A and a larger Trails effect for the HMD-CTT vs. the pencil-and-paper CTT. Group X Format X Trails interaction was not statistically significant (p=.080).

As participants in the OLD group had significantly fewer years of education compared with participants in the YA and MA groups (p≤.001; see *Methods*), we repeated the analysis entering years of education as a covariate. The results obtained did not change appreciably (see *supplementary material file #1*).

Regarding prevalence of errors, more errors occurred during performance of the HMD-CTT (Figure 6, p≤.004). Interestingly, significant group effect was found only for HMD-CTT (Trails A and B; H≥8.8, p≤.012) but not for the pencil-and-paper CTT format (H≤2.96; p≥.227). Post-hoc analyses reveal that the source of the significance is the higher number of errors among the OLD as compared to the YA (p=.009).

#### Correlations between Pencil-and-Paper and HMD-CTT - completion times

Figure 7 shows the relationship between performance on the pencil-and-paper and HMD-CTT for the YA (blue), MA (orange) and OLD (green) group. The Spearman correlation (rho; *r*_*s*_) between Part A completion time (t_A_) on the gold-standard pencil-and-paper CTT and the corresponding Part A completion time on the HMD-CTT was 0.62 (*p*<.001; Fig. 7A). For Part B completion time (t_B_), the Spearman correlation was 0.69 (*p*<.001; Fig.7B).

#### Qualitative analysis of manual performance

Figure 8 shows grand averages of the scaled ball-to-ball hand trajectory velocity profiles (see methodologies in *supplementary file # 1*) for all participants who completed HMD-CTT Trails A (solid traces) and B (dashed traces). Data from the three age groups are color-coded (see legend for details). Based on these traces, the following observations can be made:

1. For all groups and for both Trails A and B, movement toward a target (ball) does not stop immediately upon reaching (virtually touching, at x=100% which for all, except for the last trajectory, is also x=0% of the next trajectory) the target, but somewhat later, as reflected by the ensuing gradual decrease in velocity decrease at x=0%positive deflection and the ensuing gradual decrease in velocity. This decrease on the grand averaged traces does not reach zero because the minimum is reached at a different time for each of the 24, individual ball-to-ball trajectories (see *Fig. S-1* in *supplementary file #1*).
2. In Trails A, but not in Trails B, soon after reaching this minimum (i.e., completing the execution of the previous trajectory), a new trajectory can be identified (time of emergence is designated by the left black arrow). The velocity profile of this trajectory is characterized by an accelerating portion (peak indicated by a gray arrow) and a decelerating portion upon approaching the target ball. The degree of asymmetry between these portions of the trajectory varies between cohorts, with the YA showing greater symmetry.
3. Conversely, in Trails B, a prolonged ‘executive function’ period is evident, and acceleration towards the target is identifiable after at least 40% of the trace, with the older groups showing a more delayed start (black arrows on dashed traces). This pattern is consistent with the divided attention aspect of Trails B in which the participant must locate the ball with the next number in the sequence of opposite color to the current ball, ignoring the distracter ball will the correct number but the incorrect color (i.e., same color as the current ball).
4. Consistent with the results on completion times (t_A_, t_B_; Table 2), the velocity profiles for Trails A are faster than those of Trails B.

**Figure 8:**
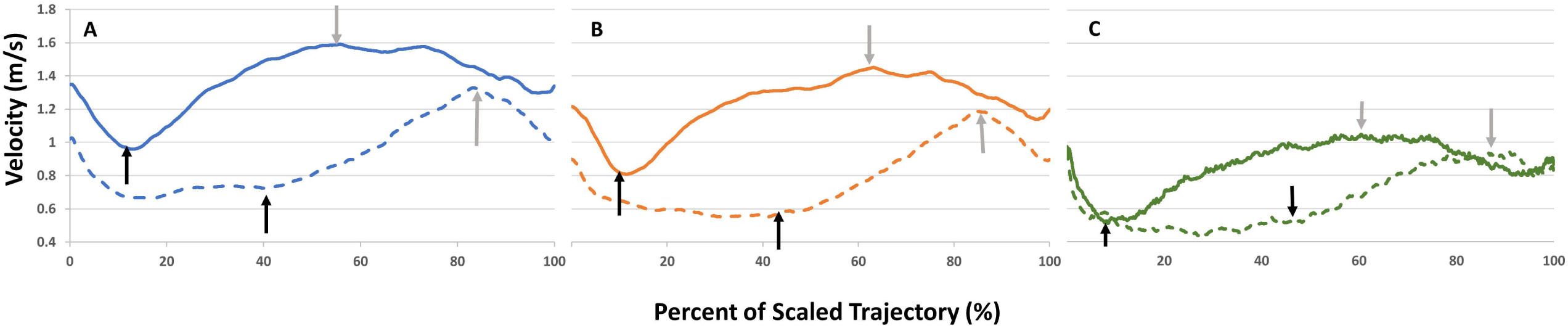
Grand averages of HMD-CTT hand movement velocity profiles. HMD-CTT hand movement velocity (meters/second) profiles showing trajectories for young (panel A), middle-aged (panel B), and older adult (panel C) groups for Trails A (solid line) and Trails B (dashed lines) over time, scaled to reflect percent from trajectory completion duration. In panel A (YA), for each of Trails A and B, 840 trajectories were avaraged from 35 participants; in panel B (MA),1176 trajectories were averaged from 49 participans; in panel C (OLD), 336 trajectories were averaged from 14 participants. See *text* for additional description and interpretation.

**Table 1:** Mean values for CTT completion times (mean ± SD)

|  | **Pencil-and-Paper CTT** | **DOME-CTT** |
| --- | --- | --- |
| YA: $t_A$ (s) | 31.01 ± 7.62 | 137.39 ± 23.84 |
| YA: $t_B$ (s) | 61.09 ± 14.66 | 213.16 ± 53.18 |
| MA: $t_A$ (s) | 50.81 ± 16.80 | 219.87 ± 67.31 |
| MA: $t_B$ (s) | 86.53 ± 21.22 | 311.26 ± 98.61 |

YA – Young adults; MA- Middle Aged; s – seconds; $t_A$ – completion time of Trails A; $t_B$ – completion time of Trails B;

**Table 2:** Mean values for CTT completion times (mean ± SD)

|  | **Pencil-and-Paper CTT** | **HMD-CTT** |
| --- | --- | --- |
| YA: $t_A$ (s) | 29.78 ± 9.08 | 69.93 ± 17.03 |
| YA: $t_B$ (s) | 61.52 ± 20.53 | 119.82 ± 25.23 |
| MA: $t_A$ (s) | 40.15 ± 15.02 | 91.36 ± 32.60 |
| MA: $t_B$ (s) | 82.60 ± 27.88 | 152.24 ± 47.99 |
| OLD: $t_A$ (s) | 63.55 ± 35.95 | 126.75 ± 84.26 |
| OLD: $t_B$ (s) | 126.57 ± 45.92 | 236.52 ± 88.00 |

YA – Young adults; MA- Middle Aged; EA – Elderly adults; s – seconds; $t_A$ – completion time of Trails A; $t_B$ – completion time of Trails B;

### Test-retest reliability (Studies 1 & 2)

For the DOME-CTT (retest interval of about 12 weeks), moderate reliability was found: Trails A (t_A_) and B (t_B_) (ICC=.597, p=.023, ICC=.676, p=.018, respectively). For the pencil-and-paper CTT, comparatively better reliability was found for Trails A (t_A_) (ICC=.778, p=.001) and for Trails B (t_B_) (ICC=.622, p=.040).

For the HMD-CTT (retest interval of about 12 weeks), moderate reliability was found for Trails A (t_A_) and B (t_B_) (ICC=.618, p=.004; ICC=.593, p=.003, respectively). For the pencil-and-paper CTT, comparatively better reliability was found for Trails A (t_A_) and Trails B (t_B_) (ICC=.744, p<.001; ICC=.769, p<.001, respectively).

For the HMD-CTT (retest interval of about 2 weeks), good reliability was found for Trails A (t_A_) and B (t_B_) (ICC=.748, p=.002; ICC= .893, p<.001, respectively). The paper-and-pencil CTT also showed good reliability for both Trails A (t_A_) and Trails B (t_B_); Compared to HMD-CTT, the gold standard test had higher reliability for t_A_ and lower reliability for t_B_ (ICC=.851, p<.001; ICC=.798, p=.001, respectively).

### Discriminant Validity (Studies 1 & 2)

Table 3 shows AUC values from ROC curves (depicted in the *supplementary file #1*). Pencil-and-paper CTT AUC values are compared to the AUC values obtained from the performance on each of the VR adaptations (i.e., DOME-CTT or HMD-CTT). For each VR adaptation, the relative difference [%] in AUC values from the pencil-and-paper AUC values version is shown. Comparisons were made separately for Trails A and Trails B.

**Table 3:**
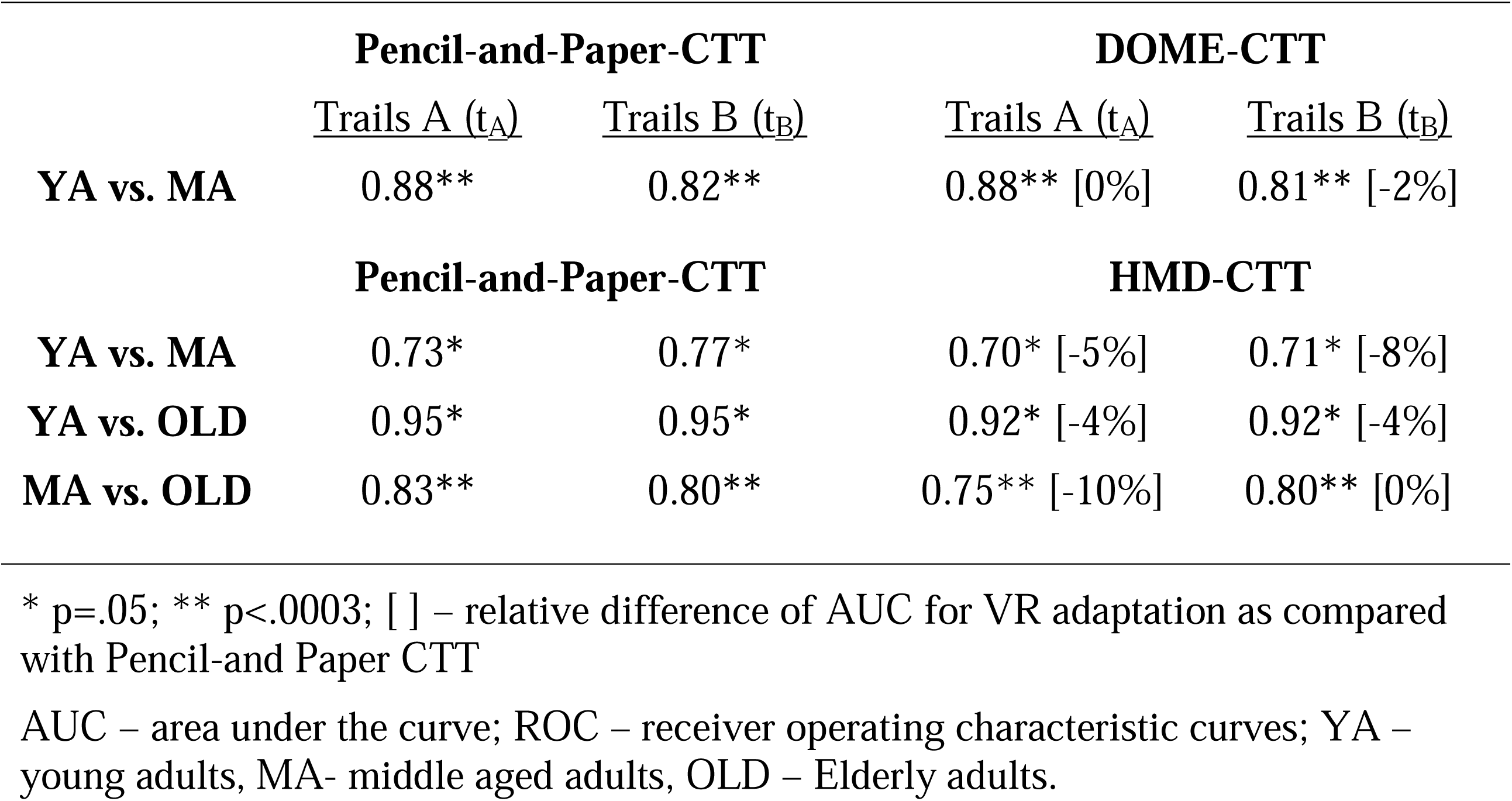
AUC values from ROC curves

The data indicate that discriminant validity of all versions of the CTT is relatively high (AUC≥ .70; p≤.05). AUCs are largely comparable, though slightly lower for the VR adaptations.

#### Comparing completion times between DOME-CTT and HMD-CTT

The completion times in Tables 1 and 2 suggest that t_A_ and t_B_ are higher (i.e., longer) for DOME-CTT relative to HMD-CTT. As none of the participants completed both DOME-CTT and HMD-CTT testing, the present work was not designed to compare between the two VR platforms. However, we conducted analyses to address this question based on the existing evidence. The methodology and the results which are detailed in the supplemental material (*supplementary file* #1) do suggest that shorter completion times for the HMD-CTT as compared to the DOME-CTT (Tables 1 and 2).

## DISCUSSION

In this study, we describe the development and initial validation of VR-based adaptations of the traditional pencil-and-paper Color Trails Test (CTT) (27) test of attention and processing speed using two completely different VR systems. The first VR-based system involves projection of visual stimuli on the walls of a large, dome-shaped room (akin to a cave). The second VR-based system is a low-cost head-mount VR device (HMD) worn by the participant and suitable for home-based assessment. Both VR-based adaptations proved to be relatively acceptable, with only two participants (∼1.5%) not completing the VR tasks. Participants only rarely complained about the difficulty of completing the VR tasks, but there were no such complaints for the pencil-and-paper version. Our discussion integrates the results from the two studies.

### Construct validity

Our results suggest that the new VR-based adaptations and gold standard pencil-and-paper version share similar psychometric properties (e.g., longer completion time for B vs. A). Coupled with the relatively high correlations between corresponding parts (∼0.7; Figs. 5&7), this suggests that the VR and the pencil-and-paper tests measure the same cognitive constructs (e.g., sustained, divided attention). By comparison, in a cross-validation study of the TMT and CTT, Dugbartey and colleagues reported lower correlation values of .35 for Trails A and .45 for Trails B (35).

Notably, construct validity correlations for YA participants on the Trails A portion of the test were not significant. We attribute this result to performance to a ceiling effect in that t_A_ values were close to the shortest completion times technically possible on all three CTT tests.

### Completion-time format effects

Trails A and Trails B completion times were significantly longer for the VR-based adaptations compared with the pencil-and-paper CTT, possibly reflecting a larger dynamic range of performance for the VR versions (e.g., even only due to the larger space covered by the hands to perform the VR task as compared to one page distribution of the targets in the pencil-and-paper CTT), greater task difficulty (36) and/or that the VR versions incorporate cognitive-motor interactions not measured by the pencil-and-paper version.

Among the two VR adaptations, completion times for the DOME-CTT were longer than those for the HMD-CTT (compare Tables 1&2, Figs. 5&7; post-hoc analyses comparing across studies – *supplementary file #1*). One possible account for this finding relates to different levels of visual immersion between the tests. The HMD-CTT provides no visual feedback from the arms, and the participant’s subjective experience of the task consists solely of moving the avatar (red ball) within the VR environment. In contrast, during the DOME-CTT, the participant sees his/her hand holding the wand-like stick in addition to the virtual avatar as s/he makes reaching movements toward the target balls. The latter configuration may complicate sensorimotor integration given the two parallel, relevant sensory input streams (physical hand, virtual avatar); this hypothesis should be further investigated.

### Error performance

Participants made significantly more errors (i.e., touching the wrong ball) on the VR-based versions as compared to the pencil-and-paper CTT (Figs. 4&6). For example, only about 14% of all pencil-and-paper test levels completed in Study 2 (Trials A and B across all three cohorts) had at least one error (most often one error). This error rate was 2.5 times higher for HMD-CTT test levels (∼35%).

This pattern of results may seem paradoxical as with longer completion times on the VR-based tests, fewer errors should occur, but our data reflect the opposite. Indeed we believe that as the VR tasks are more demanding then the corresponding pencil-and-paper tasks, the cognitive processes classically associated with the CTT paradigm might be compromised, leading to more errors (37-40). Specifically, we posit that the VR-based tasks make much greater demands on motor planning and execution (see below), visual scanning and spatial orientation and involve higher perceptual and/or cognitive load. Apparently, this load differentially affect elderly as compared to YA as evident by the significant higher error rates recorded among the OLD group.

### Cognitive-motor interactions

Our qualitative data clearly demonstrate that when shifting from a cognitive task primarily involving sustained attention (Trails A) to one that primarily involves divided attention (Trails B), associated upper-limb motor behavior changes (Fig. 8). Previous research has employed VR to study cognitive-motor interactions mainly in the context of locomotion. Most of the studies have reported clinical benefits related to cognitive-motor interactions associated with immersion in a VR environment (41-45).

In this study, we began exploring the effect of divided attention (as operationalized by HMD-CTT Trails B performance) on the planning and execution of manual (upper-limb) reaching movements. The well-documented single-peak velocity profile typical of ballistic movements (46-48) appears to govern the hand trajectories generated during HMD-CTT Trails A. However, this is not the case for Trails B, for which trajectories are characterized by an initial slow increase in the velocity profile, probably reflecting neural resources more engaged in executive function and less in motor execution during a divided attention task. Potential age effects (e.g., less symmetric peak, slower overall velocity) are apparent in comparing the velocity profile across age groups (Fig. 8).

Follow up studies will focus on developing reliable quantification methods and metrics to assess these cognitive-motor interaction effects. Notably, comparing the velocity profiles generated during a three-dimensional (3D) VR-based task to a classical two-dimensional (2D) task as the ‘gold standard’ (49) is suboptimal, mainly due to the absence of a reliable theoretical model for three-dimensional hand-reaching movements. Thus, new referencing methodologies like sampling single target-to-target single trajectories should be included as part of future versions and analyses of VR-based CTT tasks like those used here.

### Discriminant validity

The VR-based tests were largely comparable though not superior to the pencil-and-paper CTT in terms of distinguishing individuals of different age-groups on the basis of CTT completion time (t_A_ and t_B_, Table 3). Further, both the traditional and VR-based versions demonstrated relatively high discriminant validity as reflected by high AUC values. These observations are consistent with the strong correlations for completion times between the VR-based and original CTT for each age group (with the exception of t_A_ in YA) as well as when combining participants across age groups (black dashed lines in Figs. 5&7).

Comparability in discriminant validity between VR-based and the gold standard CTT for completion times is encouraging. However, VR-based testing affords additional metrics that may be better at differentiating among age groups and among healthy and cognitively impaired individuals. Indeed, VR facilitates the development of new, more ecologically valid parameters more relevant to daily living in that they better capture more complex, integrated behaviors. Thus, we speculate that using such VR-based parameterization of multimodal function (e.g., hand-gaze coordination combined with hand trajectories) will provide superior discriminant validity.

### Test retest reliability

For a retest period of ∼12 weeks, the VR-based CTT adaptations showed moderate reliability (intraclass correlation of ∼0.6), while the pencil-and-paper version generally showed comparatively better reliability. The superior reliability of the original CTT for this retest interval may be attributable to the greater familiarity of the pencil-and-paper format as compared to the VR, which may have led to better performance (i.e., a learning effect) upon retest and consequently poorer reliability (see (50)). We also acknowledge that some members in the reference cohort had undergone a cognitive training protocol during the 12-week interval (see *table S1* in the *supplementary material*) which may compromise test-retest evaluations.

However, for a retest period of ∼2 weeks, both the HMD-CTT and the original CTT showed good reliability (intraclass correlation of ≥0.75), with the VR-based adaptation showing superior reliability for Trails B. Our findings are consistent with findings in the literature that shorter retest intervals yield higher reliability coefficients (51). Notably, there does not appear to be a clear convention or recommendation for the ideal retest interval (52). Thus, our data reporting reasonable reliability for two different retest intervals is relevant and informative. It should be noted, however

Still, as sample sizes were small given that only a subset of our participants had data for each retest interval, it would be good to replicate our findings in larger samples.

### Future Directions

VR technologies may enable us to enrich the current VR-based versions of the CTT to further enhance ecological relevance, mainly in the sense of engaging more modalities, and inter-modalities interactions.

The challenge will then be how to leverage multimodal measures to understand such real-world processes as cognitive-motor interference during multi-tasking and ultimately assess function in a predictive or clinically meaningful way. In particular, we hope to achieve superior discriminant validity for patient cohorts and the ability to predict risks associated with impaired cognitive-motor interactions (53-56), such as the risk of falls in the elderly and in neurological patients (57, 58).

Finally, we envision developing adaptations of additional neuropsychological tests, with different core construct (i.e., than the CTT) for application in an immersive VR environment.

### Conclusions

In sum, the present study describes the development and validation of large-scale (DOME-CTT) and head-mount (HMD-CTT) VR adaptations of the classic pencil-and-paper Color Trails Test (CTT) and provides key validation data, including construct validity relative to the original test, discriminant validity among age groups, and test-retest reliability at two different retest intervals. Critically, this work demonstrates the feasibility and viability of converting a neuropsychological test from two-dimensional pencil-and-paper to three-dimensional VR based format while preserving core features of the task and assessing the same cognitive functions. Our novel findings on the relationship between classical cognitive performance and upper-limb motor planning and execution may lead to new analysis methods for other more ecological VR-based neuropsychological tests that incorporate cognitive-motor interactions.

## List of abbreviations

ANOVA: analysis of variance
AUC: area under the curve
CTT: color trails test
FOV: field of view
HMD: head mounted device
ICC: intraclass correlation coefficients
MA: middle-aged adults
MET: multiple errands test
ROC: receiver operating characteristic
SD: standard deviation
TMT: trail making test
VET: virtual errands test
VR: virtual reality
YA: young adults

## Acknowledgments

The authors wish to thank Ms. Lotem Kribus Shmiel and Mr. Evyatar Arad for assistance in designing the DOME-CTT, Prof. Yuval Heled, Dr. Ran Yanovich, Ms. Hadar Sharon, Ms. Anastsaia Tkachov, Mr Hani Baransi Mr. Yuval Ketko, for their collaboration in data collection, and Ms. Hila Iny for technical assistance.

