## supplementary file #1 for "Multimodal immersive trail making – virtual reality paradigm to study cognitive-motor interactions"

**A. Supporting material for the *Methods* section. Subsection: *General***

**Table S1: Participants by partition to the study protocols used for collecting data for the current project**

| **Title of research** | **YA – Study 1** | **MA – Study 1** | **YA- Study 2** | **MA – Study 2** | **OLD – Study 2** |
| --- | --- | --- | --- | --- | --- |
| *Cognitive-motor rehabilitation program using virtual reality for middle-aged individuals at high dementia risk* [1] | - | 29 | - | 48 | - |
| **CTT testing procedure:** As part of the larger interventional protocol, the participants completed cognitive assessments pre-, post- and at three-month follow up after target or control intervention (application of the intervention lasted 12 weeks). CTT data included in the present report were from the pre- assessment. Administration order of VR-based CTT and pencil-and-paper CTT was counterbalanced and performed on the same day. Data from the second (post-) visit were included for test-retest reliability analyses (see text). | | | | | |
| *Characterization of cognitive-motor performance in virtual reality-based cognitive assessments.* | 10 | 0 | 36 | 3 | 17 |
| **CTT testing procedure:** As part of the larger cross-sectional protocol, the participants were assessed at two time points (two weeks apart). Exceptions to this were the 10 YA included in Study 1 who came only for one visit. During each visit, the participants were assessed with a battery of cognitive tests, including pencil-and-paper CTT and VR-based CTT, administered in counterbalanced order. In the manuscript, data collected from the first visit are included. Data from the second visit were included for test-retest reliability analyses (see text). | | | | | |
| *Evaluation of physical and cognitive performance after simulated road march combines physical and cognitive load using a virtual reality* [2] | 4 | - | - | - | - |
| **Description of CTT testing:** As part of the larger cross-sectional protocol, the participants were assessed at three time points (about one week apart). During each visit, they were assessed with cognitive tests twice: before and after one of the following three exposures, randomly presented: (1) physical load applied during 2 h treadmill march in large-scale VR facility (see Fig. 1 in the main paper) with 30% body weight load carried; (2) Same as (1) with additional cognitive load; (3) rest. CTT data included in the present report were from the first time point during the pre-exposure assessment. Administration of the pencil-and-paper CTT preceded that of the DOME-CTT. | | | | | |
| **Total:** | **YA- Study 1** | **MA – Study 1** | **YA- Study 2** | **MA – Study 2** | **OLD – Study 2** |
|  | 14 | 29 | 36 | 51 | 17 |
| YA- young healthy adults; MA – Middle-aged healthy adults; OLD – healthy older adults | | | | | |

**B. Supporting material for Study 2 Methods**

*Qualitative analysis of manual performance*

Spatial coordinates of the controller position (corresponding to the virtual ‘red ball’ avatar) were recorded throughout the HMD-CTT sessions. Custom software written in MATLAB (Mathworks, Inc.) used this data to extract and analyze the 24 target-to-target reaching movements during Trails A and Trails B, respectively. Specifically, as part of this study, we aimed to qualitatively identify and characterize different patterns in the velocity profiles of manual trajectories executed during Trails A and B. Manual (upper-limb) velocity profiles are common outcomes in studying motor control models. For example, the minimal-jerk model proposed by Flash et al. [3], predicts the hand velocity profile between two points on a plane. The predicted profile is characterized by a bell-shaped curve. Figure S-1 shows data from a typical YA participant to illustrate the procedure used to extract the velocity profiles. Velocity profiles were derived for all 24 three-dimensional ball-to-ball (target-to-target) paths (thin colored traces; Trails A – Fig. S-1A; Trails B – Fig. S-1B). Total movement time was scaled from 0 to 100% to give a scaled trajectory (Fig. S-1C&D for Trails A and B, respectively), which allows superimposing all trajectories and computing a grand average (thick black lines). In Study 2, we computed the grand average of all grand averages from individual participants to exploring the velocity profiles of manual trajectories during performance of the HMD-CTT.

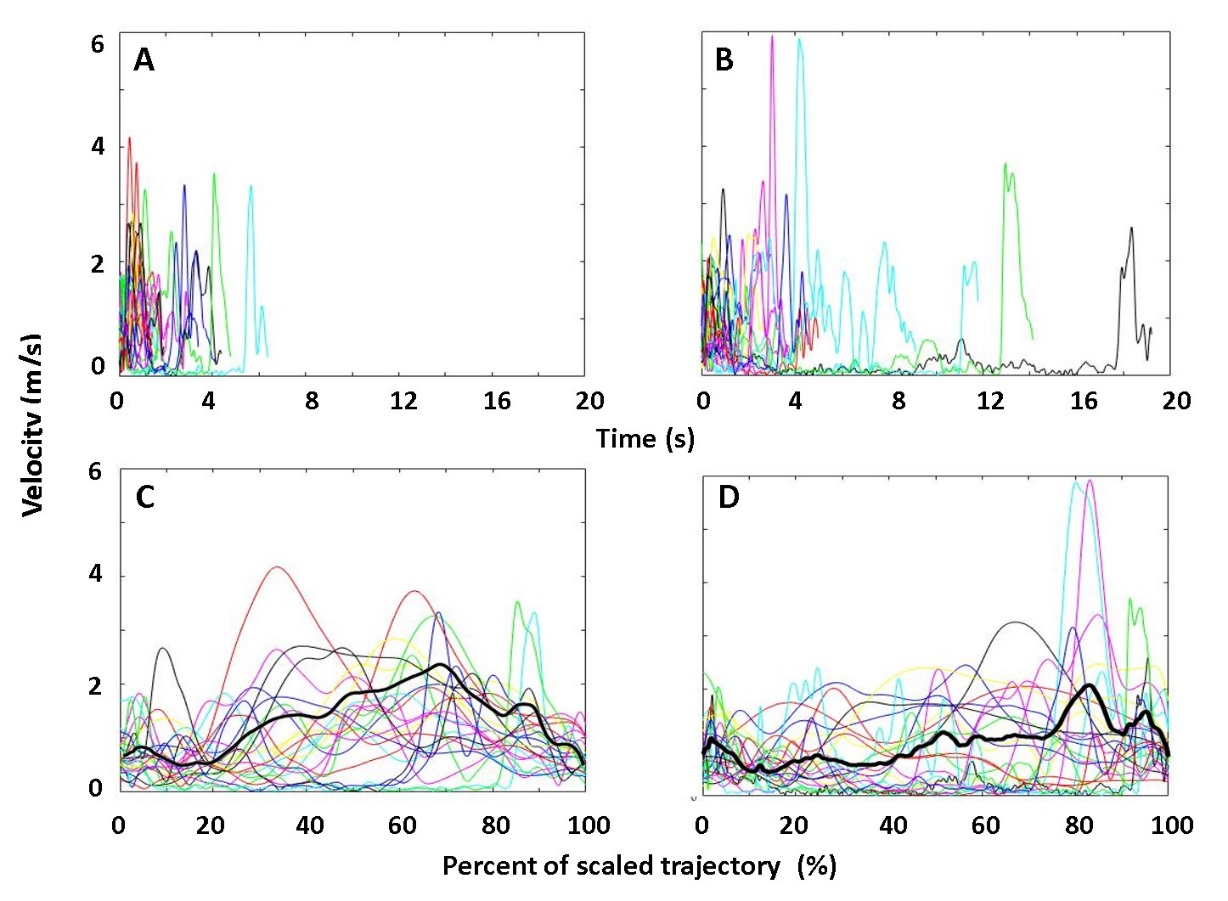

**Figure S-1: Upper-limb (manual) performance during the HMD-CTT**

In the three-dimensional HMD-CTT, the participant completes the task while standing and wearing a VR headset. S/he holds a VR controller in his/her dominant hand to move the (red ball) avatar to sequentially connect numbered balls distributed throughout the virtual space visible in the headset. Based on time series position data from the controller, velocity profiles of all target-to target paths are plotted (thin colored lines) for Trails A (panel A) and B (panel B). It is apparent that the former were generally executed in a shorter time than the latter. Next, total movement time was scaled from 0 to 100% to give a scaled trajectory. Scaled grand average trajectories were superimposed graphically (panels C and D for Trails A and B, respectively). For each percent increment, the mean value was calculated using all values of this percent in the individual traces. Averaging all mean values yielded the grand averages (thick black lines).

**C. Supporting material for the Results for study 2 section**

*Performance on the HMD-CTT: Group and Format effects –with ‘education years’ as a covariate*

Statistical analysis revealed a significant effect of Group (F_2,97_ = 34.1, p<.0001, η²=.41; progressively longer completion time with more advanced age, all pairwise comparisons p<.001) an effect of Format (F_1,97_ = 22.4, p<.0001, η²=.18; longer completion time for HMD-CTT) and an effect for Trails (F_1,97_ = 11.3, p=.001, η²=.10; longer completion time for Trail B). The Group x Format and the Trails X Group interactions were also statistically significant (F_2,97_ = 8.4, p<.0001, η²=.14, F_2,97_ = 13.0, p<.0001, η²=.21, respectively).

**D. Supporting material for the results on discriminant Validity (Studies 1 & 2)**

*Depiction of the receiver operating characteristic (ROC) curves*

Table 3 details the area under the curve (AUC) values calculated from the ROC curves. Herein the ROC curves are presented in corresponding order to the entries in Table 3.

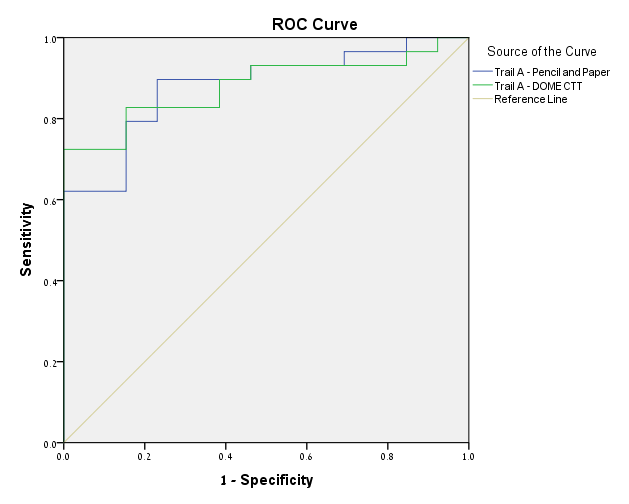

Discrimination between YA and MA – Pencil- and- Paper CTT vs DOME-CTT, Trails A

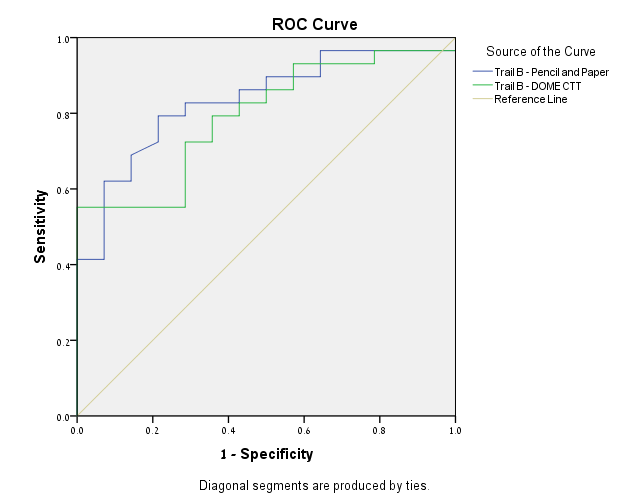

Discrimination between YA and MA – Pencil- and- Paper CTT vs DOME-CTT, Trails B

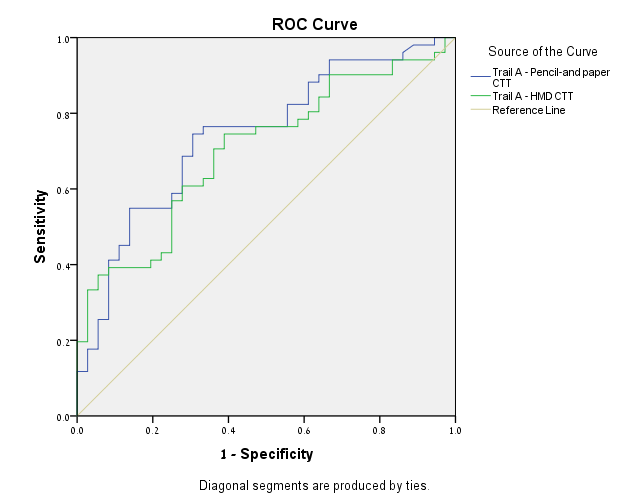

Discrimination between YA and MA – Pencil- and- Paper CTT vs HMD-CTT, Trails A

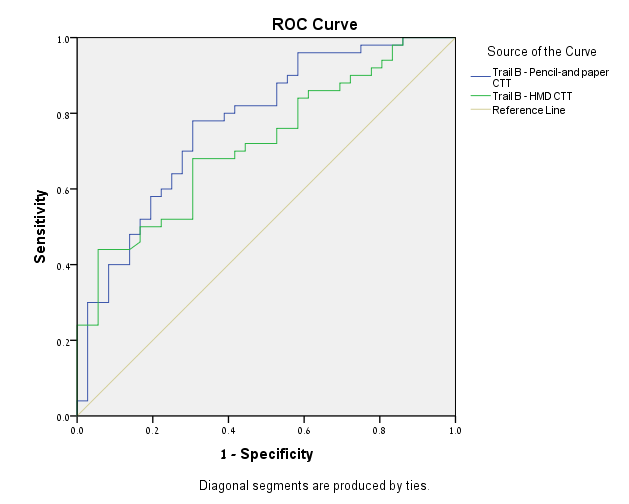

Discrimination between YA and MA – Pencil- and- Paper CTT vs HMD-CTT, Trails B

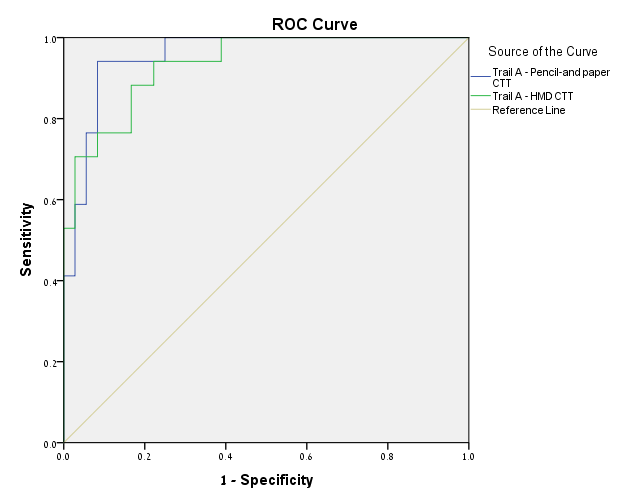

Discrimination between YA and OLD – Pencil- and- Paper CTT vs HMD-CTT, Trails A

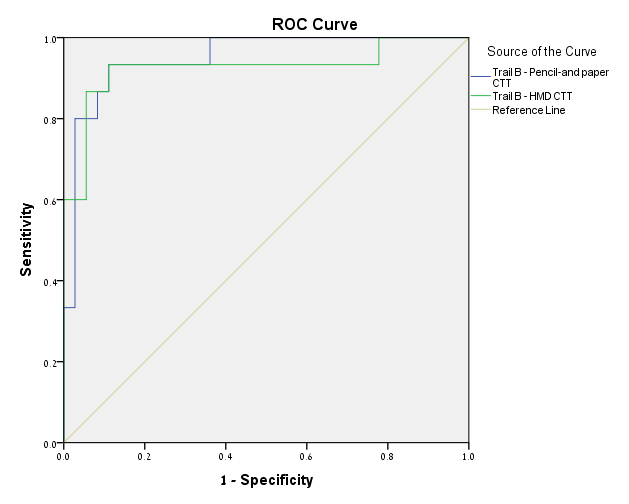

Discrimination between YA and OLD – Pencil- and- Paper CTT vs HMD-CTT, Trails B

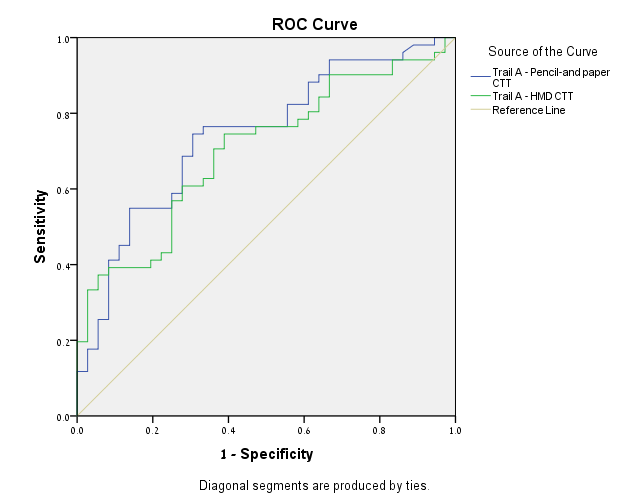

Discrimination between YA and MA – Pencil- and- Paper CTT vs HMD-CTT, Trails A

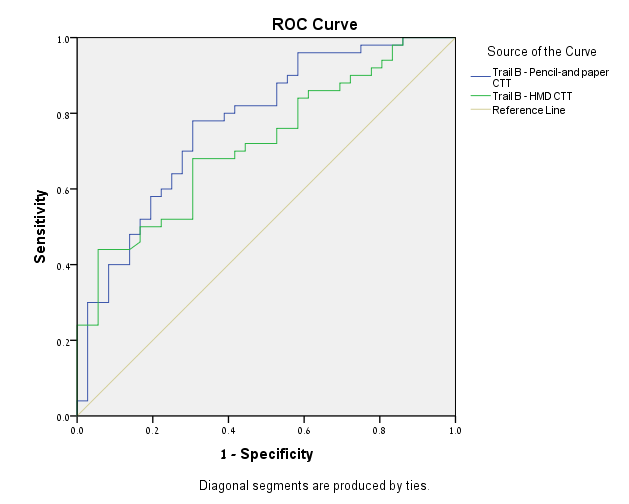

Discrimination between YA and MA – Pencil- and- Paper CTT vs HMD-CTT, Trails B

**e. Supporting material for cross study analyses**

*Comparing completion times between DOME-CTT and HMD-CTT*

The completion times in Tables 1 and 2 suggest that t_A_ and t_B_ are higher (i.e., longer) for DOME-CTT relative to HMD-CTT. As none of the participants completed both DOME-CTT and HMD-CTT testing, we adopted the following approach to compare completion times from the two VR-based adaptations (i.e., DOME-CTT vs. HMD-CTT). First, we combined paper-and-pencil CTT completion times for YA and MA participants, separately for Study 1 and Study 2. (OLD Study 2 participants were excluded because there was no Study 1 data in this age group.)

A repeated measures analysis revealed a non-significant group effect F(_1,128_) = 3.4, p=.06, reflecting a similar performance level and suggesting that YA and MA participants could similarly be combined for DOME-CTT (Study 1) and HMD-CTT (Study 2), respectively. A “repeated measures analysis” between these two groups yielded a statistically significant Format effect F(_1,126_) = 168.4, p<.0001, attributable to shorter completion times for the HMD-CTT as compared to the DOME-CTT (Tables 1 and 2).
